## Supplementary Figures for "Naturally acquired blocking human monoclonal antibodies to *Plasmodium vivax* reticulocyte binding protein 2b"

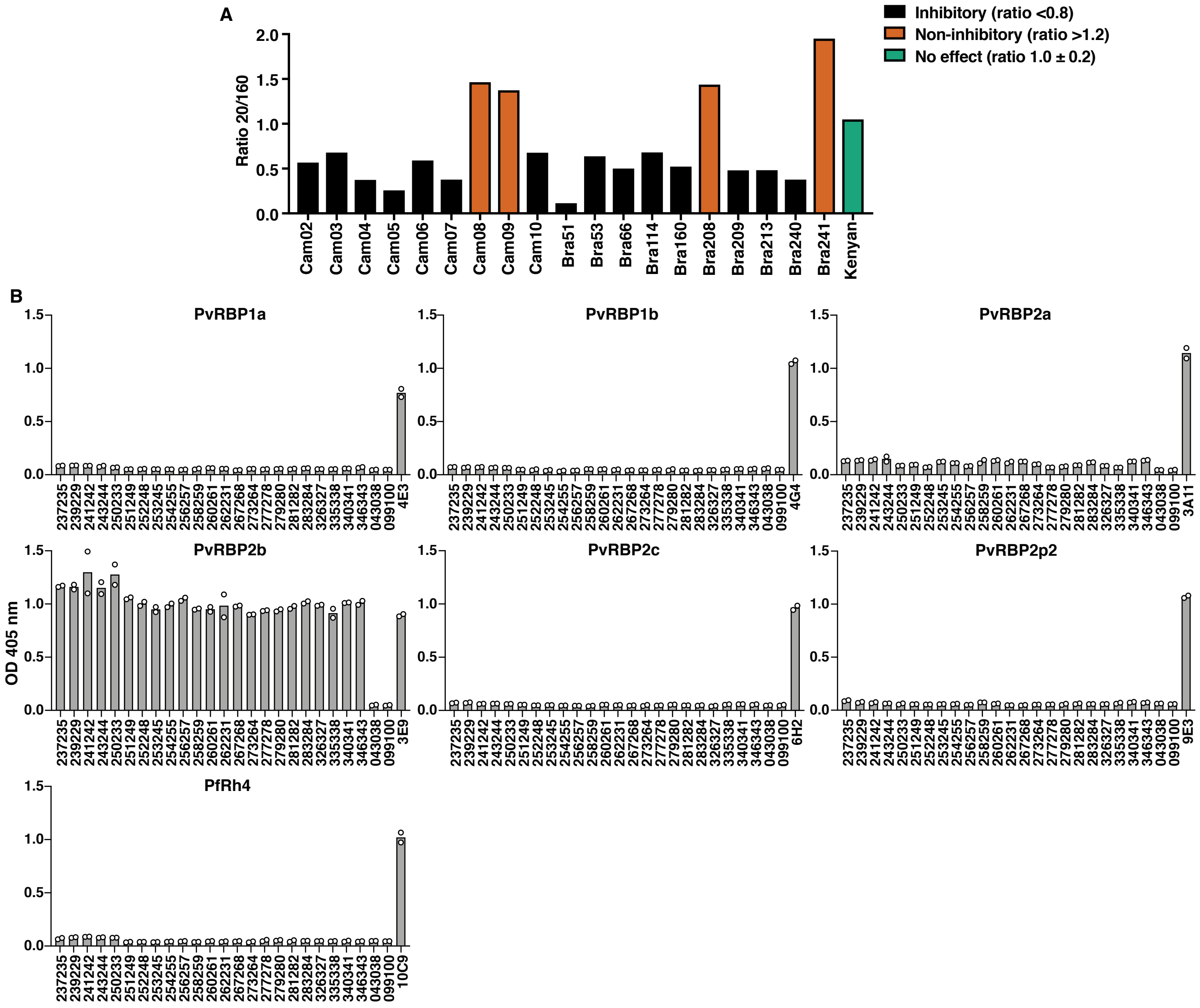


**Supplementary Figure S1. Functional blocking antibodies against PvRBP2b in serum and antibody specificity of PvRBP2b human mAbs using ELISA. (A)** PvRBP2b$_{161–1454}$ binding to reticulocytes in the presence of serum from Cambodian and Brazilian individuals and one Kenyan individual analyzed by flow cytometry. Binding results were expressed as a ratio between PvRBP2b$_{161–1454}$ binding at a 20-fold dilution of sera over a 160-fold dilution of sera, to account for non-specific increases in PvRBP2b$_{161–1454}$ binding in the presence of sera from individuals exposed to *P. vivax*. Black, ratio <0.8 and inhibitory; Orange, ratio >1.2 and non-inhibitory; Green, ratio 1.0 ± 0.2 and no effect. **(B)** Antibody specificity of PvRBP2b human mAbs to PvRBP family members and PfRh4 using ELISA. Bar graphs represent the mean of duplicate measures represented as circles. Mouse mAbs 4E3, 4G4, 3A11, 3E9, 6H2, 9E3 and 10C9 were used to detect the coating of PvRBP1a, PvRBP1b, PvRBP2a, PvRBP2b, PvRBP2c, PvRBP2p2 and PfRh4 on plates, respectively. Graphs are a representative of two independent experiments.

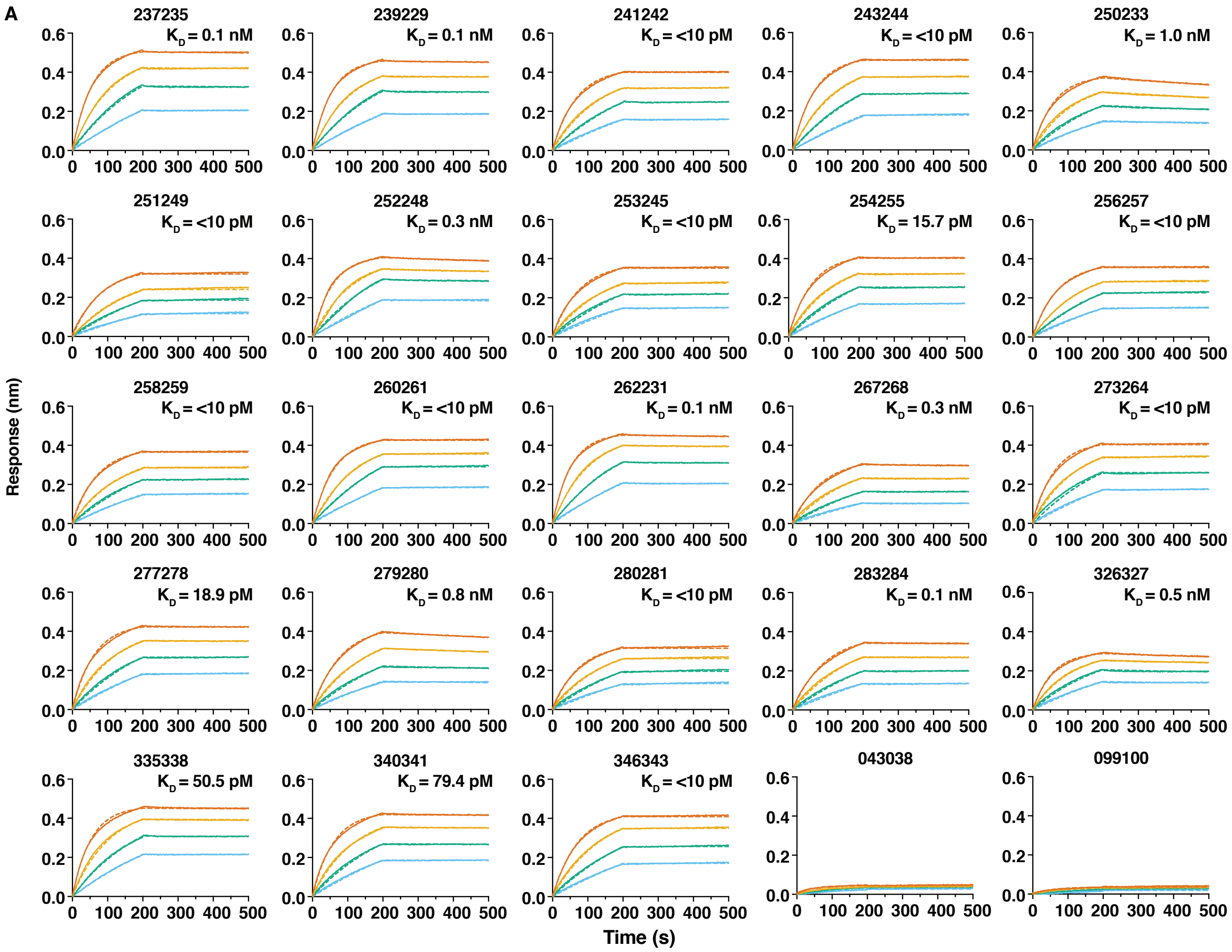


**B**

| mAb | $K_a$ (x $10^5$ $M^{-1}s^{-1}$) | $K_d$ (x $10^{-5}$ $s^{-1}$) | $K_D$ (x $10^{-11}$ M) |
|---|---|---|---|
| 237235 | 3.57 | 2.11 | 5.16 |
| 239229 | 3.80 | 2.80 | 6.58 |
| 241242 | 3.27 | 1.63 | 4.15 |
| 243244 | 3.25 | 0.73 | 0.10 |
| 250233 | 3.54 | 32.08 | 90.50 |
| 251249 | 2.75 | 0.10 | 0.10 |
| 252248 | 3.39 | 19.10 | 57.90 |
| 253245 | 2.82 | 0.64 | 1.88 |
| 254255 | 3.00 | 1.91 | 5.37 |
| 256257 | 2.86 | 0.10 | 0.10 |
| 258259 | 2.94 | 0.59 | 1.59 |
| 260261 | 3.36 | 0.49 | 1.12 |
| 262231 | 3.80 | 4.69 | 10.30 |
| 267268 | 2.31 | 4.31 | 15.80 |
| 273264 | 3.27 | 3.02 | 7.70 |
| 277278 | 3.76 | 3.08 | 7.66 |
| 279280 | 2.60 | 19.37 | 72.73 |
| 281282 | 2.63 | 0.60 | 2.13 |
| 283284 | 2.26 | 6.00 | 24.33 |
| 326327 | 3.78 | 24.57 | 64.20 |
| 335338 | 4.32 | 3.08 | 7.52 |
| 340341 | 3.37 | 3.85 | 10.41 |
| 346343 | 2.82 | 0.10 | 0.10 |

**Supplementary Figure S2. PvRBP2b human mAb binding kinetics by bio-layer interferometry. (A)** Representative sensorgrams (solid lines) and curve fitting analysis using a 1:1 model (dashed lines) for human mAbs binding to PvRBP2b$_{161-1454}$. A two-fold concentration gradient of PvRBP2b$_{161-1454}$ from 6 - 50 nM is shown by the different colored lines (50 nM, red; 25 nM, orange; 12 nM, green; 6 nM, blue). The calculated $K_D$ is shown for each sensogram. 043038 and 099100 are isotype controls. **(B)** Table of association rate constants ($k_a$), dissociation rate constants ($k_d$) and equilibrium dissociation rate constants ($K_D$) for PvRBP2b human mAbs. Table values are an average of four independent experiments.

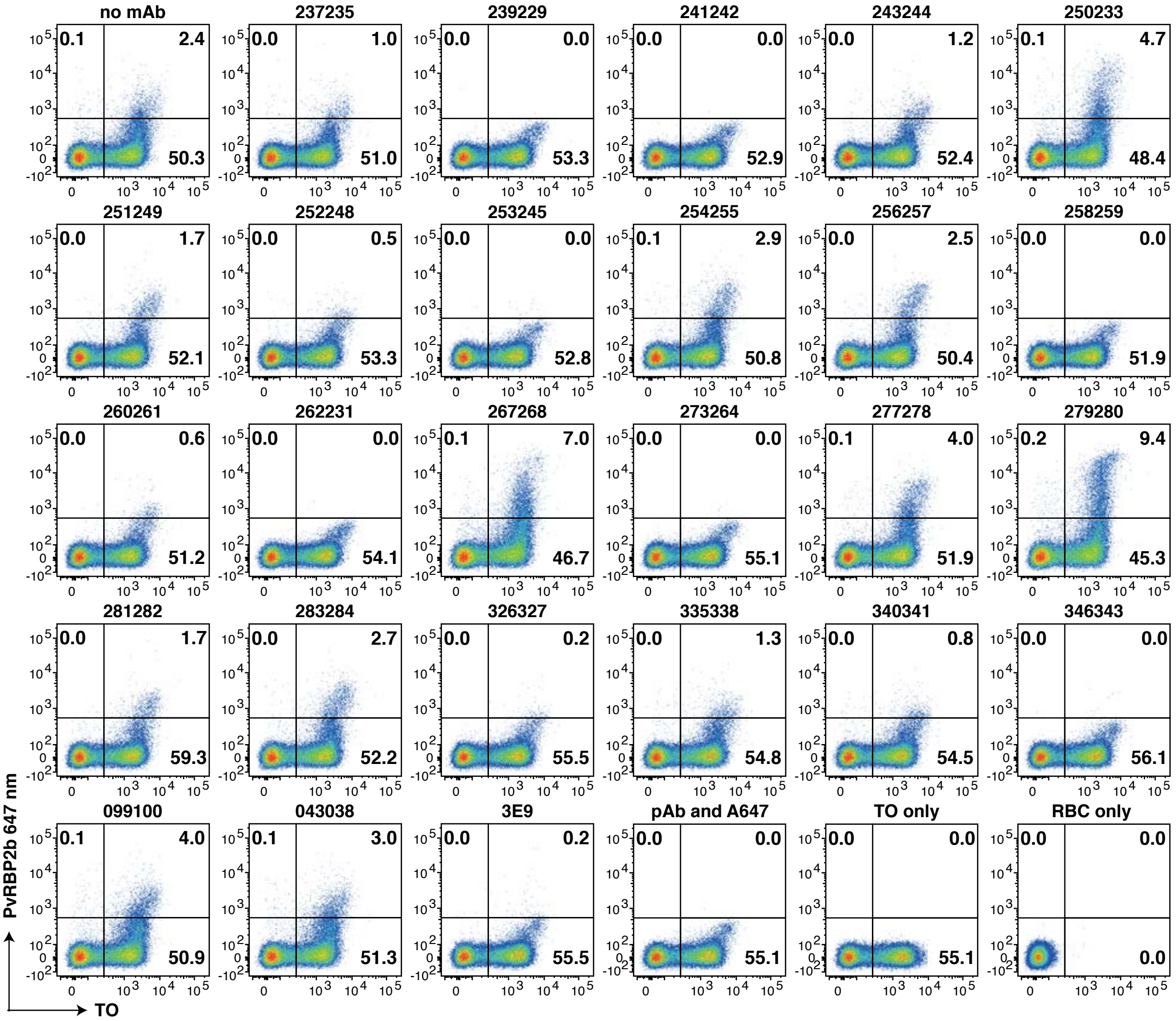


**Supplementary Figure S3. PvRBP2b human mAbs block PvRBP2b binding to reticulocytes.** Representative dot plots showing the binding of $\text{PvRBP2b}_{161-1454}$ to reticulocytes in the presence of PvRBP2b human mAbs. Thiazole orange (TO) was used to stain the reticulocyte population on the x-axis. $\text{PvRBP2b}_{161-1454}$ binding was detected with PvRBP2b polyclonal antibodies (pAb) and Alexa 647 secondary antibody on the y-axis. 099100 and 043038 were isotype controls and 3E9 was an inhibitory mouse mAb control. The reticulocyte population was gated on the red blood cell (RBC) population and the $\text{PvRBP2b}_{161-1454}$ positive population was gated on the detecting antibodies (pAb and A647) control that showed background signal to reticulocytes.

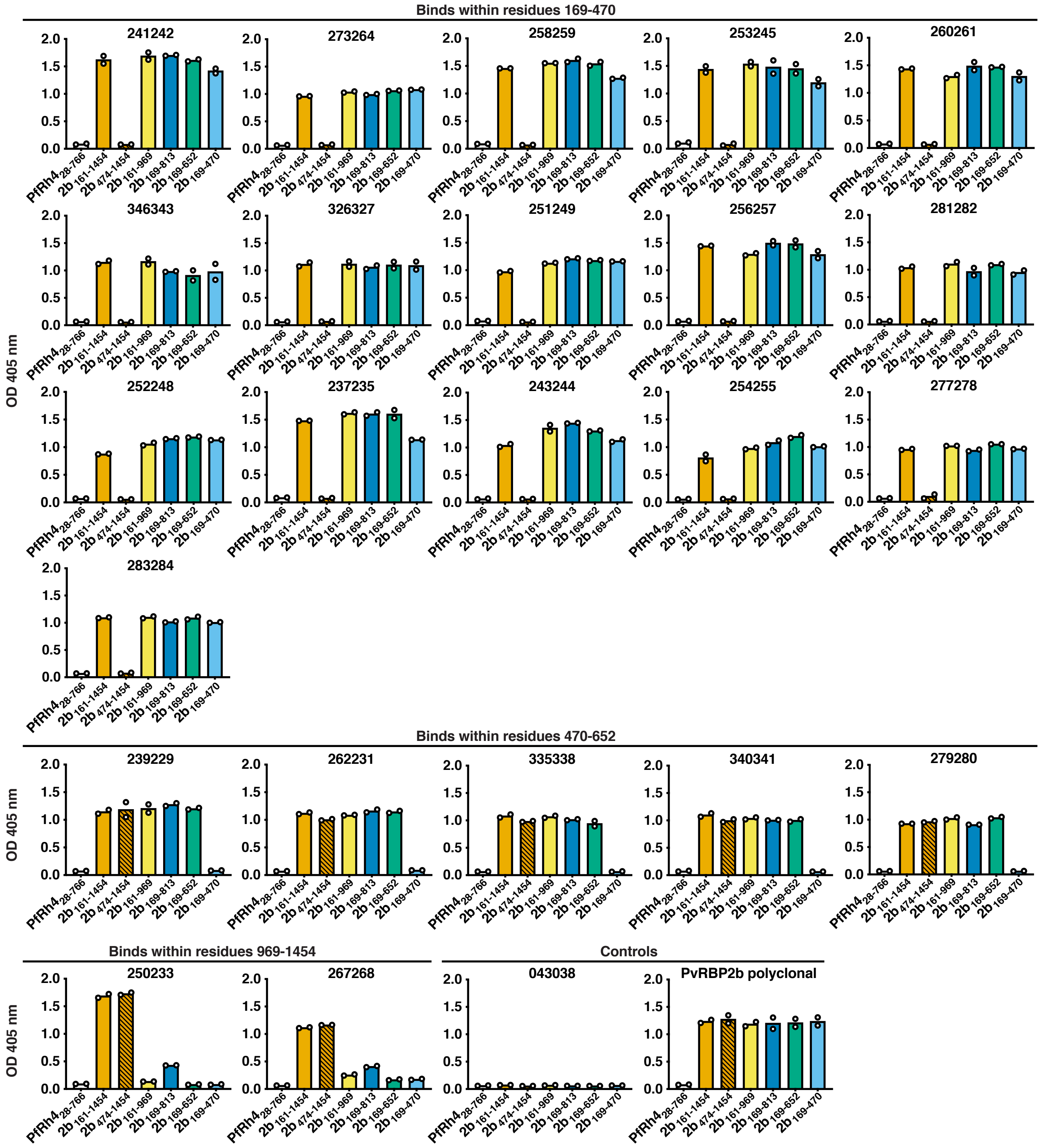


**Supplementary Figure S4. Domain mapping of PvRBP2b human mAb epitopes.** Binding of PvRBP2b human mAbs to PvRBP2b recombinant fragments and PfRh$_{428-766}$ was detected by ELISA. PfRh4$_{28-766}$ was used as a control for non-specific binding of PvRBP2b human mAbs. PvRBP2b polyclonal antibody was used to detect coating of PvRBP2b fragments and 043038 was used as a negative control. Bar graphs represent the mean of duplicate measures represented as circles. Graphs are a representative of two independent experiments.

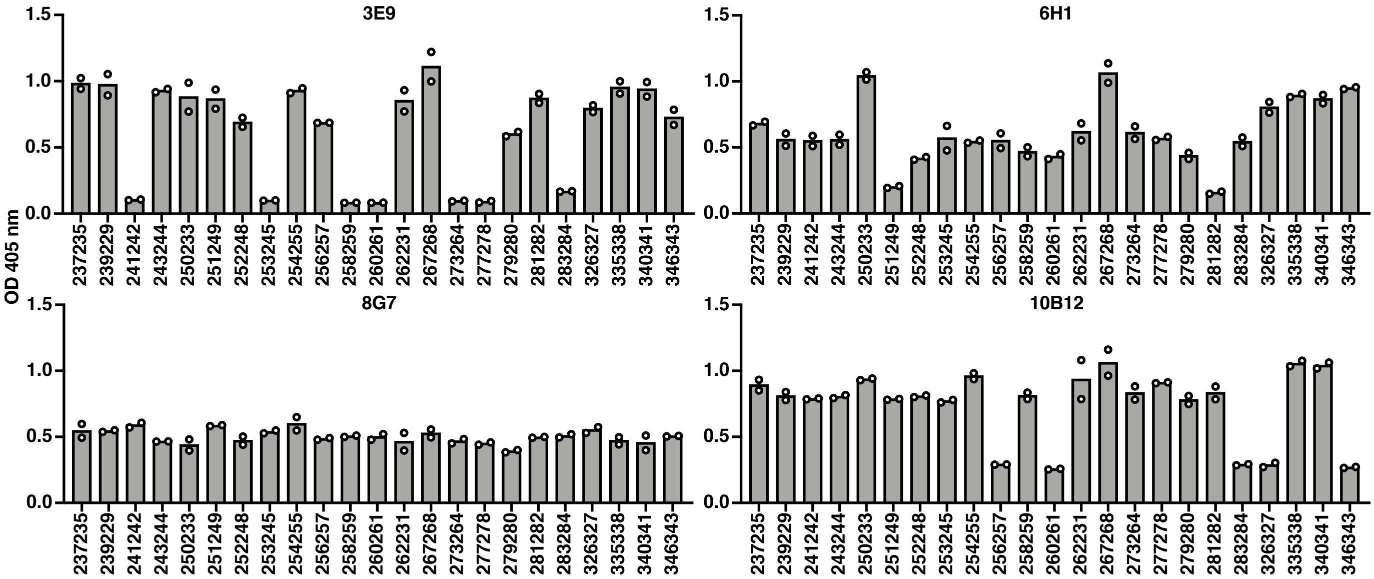


**Supplementary Figure S5. Competition between PvRBP2b human mAbs with PvRBP2b mouse mAbs.** Competition ELISA using immobilized PvRBP2b mouse mAbs incubated with a mixture of PvRBP2b human mAbs with $PvRBP2b_{161-1454}$ at a 20:1 molar ratio. Bar graphs represent the mean of duplicate measures represented as circles. Graphs are a representative of two independent experiments.

Sequence coverage: 95%

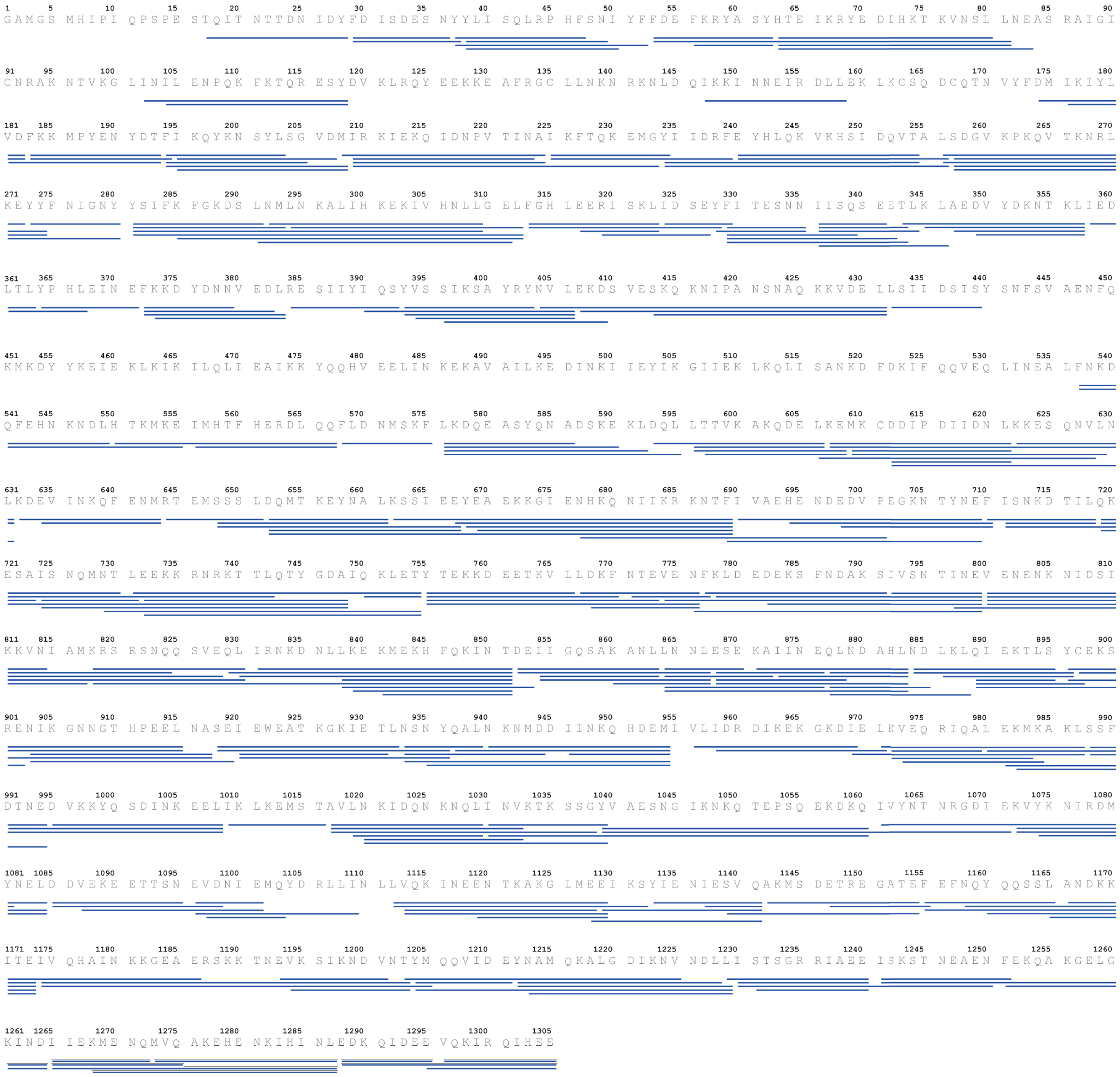


**Supplementary Figure S6. HDX peptide map for PvRBP2b$_{161\text{-}1454}$.** PvRBP2b$_{161\text{–}1454}$ peptide map showing the peptides used for HDX-MS analysis. The residue numbering corresponds to the recombinant protein and 149 residues must be added to match with the endogenous protein.

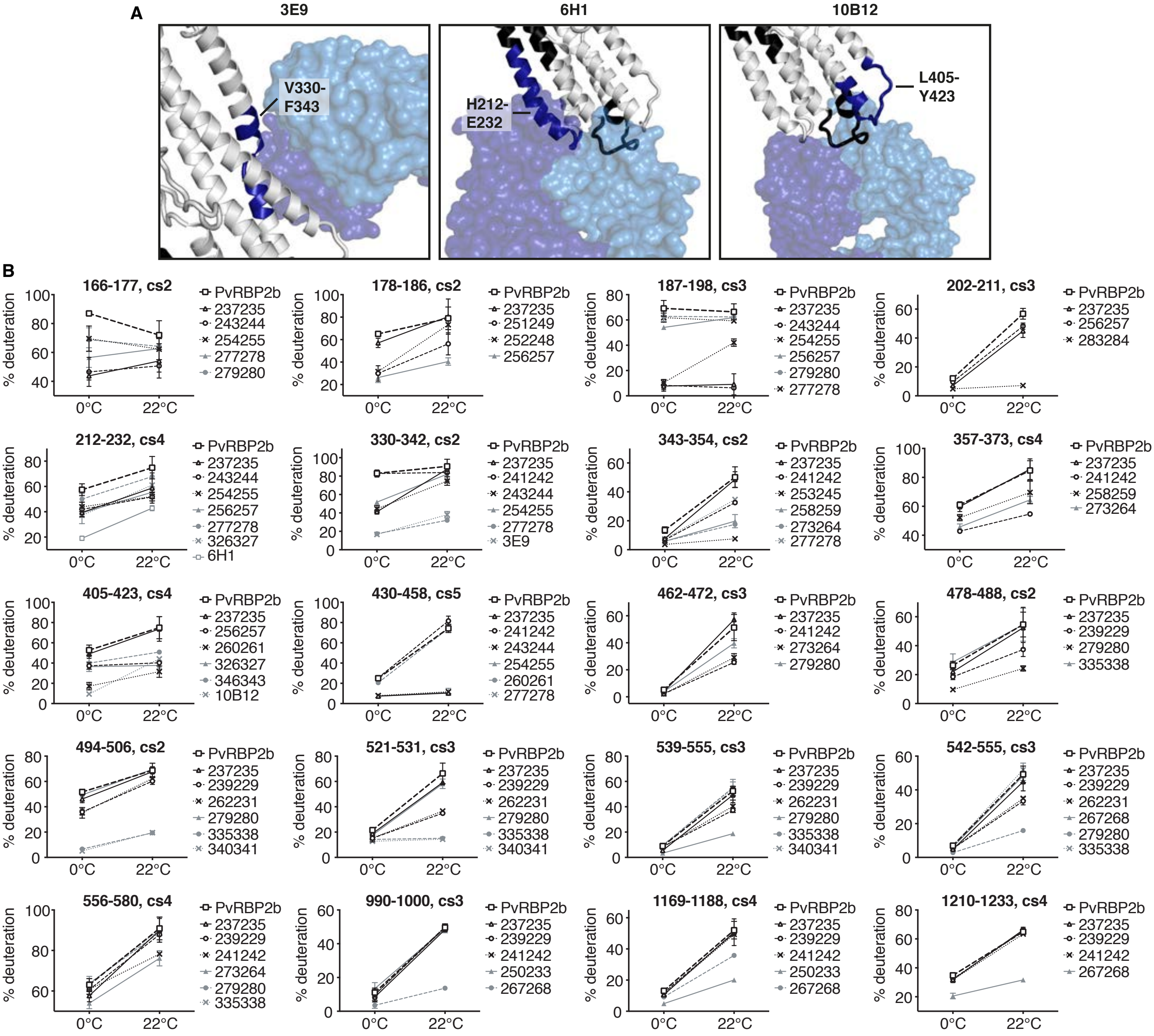


**Supplementary Figure S7. Comparison between PvRPB2b epitopes identified by HDX-MS and X-ray crystallography for mouse mAbs and HDX-MS uptake plots for a selection of PvRBP2b peptides. (A)** Combination of HDX-MS data with crystal structures of PvRBP2b mouse Fab fragments bound to PvRBP2b. Fabs shown in transparent surface representation. Regions that show protection by HDX-MS for each mAb are colored in blue and the residue range is labelled. Black indicates the regions where no peptides were detected by HDX-MS. **(B)** Uptake plots showing deuterium incorporation levels for a selection of PvRBP2b peptides. For each peptide, the level of deuteration, expressed as percentage compared to a fully-deuterated sample, is shown for samples incubated 5 min in deuterated buffer at 0°C and 22°C. One peptide representative of each of the PvRBP2b region showing protection in Fig. 4A is shown. Deuteration level of PvRBP2b alone is shown for every peptide. A selection of mAb showing either no change or a significant difference in deuteration level compared to PvRBP2b alone is shown for every peptide.

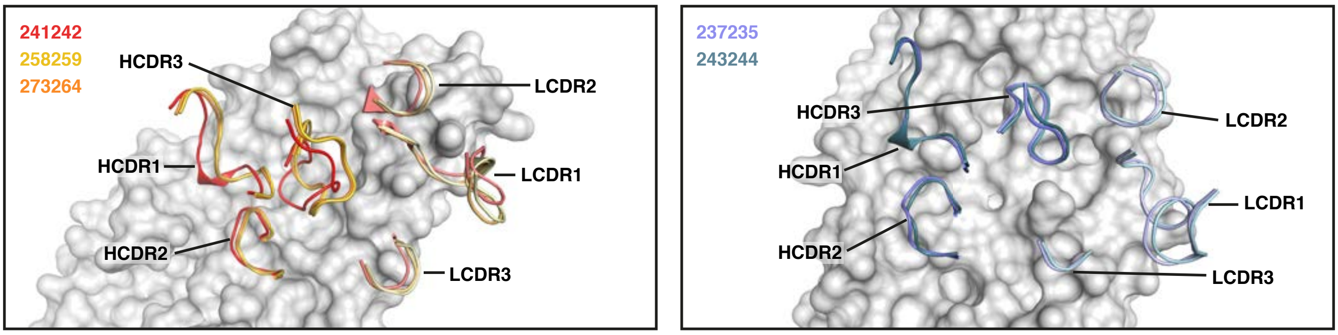


**Supplementary Figure S8. Superimposed CDR loops of antibodies with overlapping binding sites.** (Left panel) Superimposed CDR loop structures (HCDR; heavy chain CDR loops, LCDR; light chain CDR loops) of 241242, 258259 and 273264 (same clonal group as 258259) binding to PvRBP2b. (Right panel) Superimposed CDR loops of 237235 and 243244 (both from the same clonal group) binding to PvRBP2b.

**Supplementary Table S1. Diversity of VDJ gene usage for PvRBP2b human mAbs**

**Supplementary Table S2. HDX data table showing for each peptide analyzed and for PvRBP2b alone or in complex with PvRBP2b human mAbs: i) deuteration levels expressed as number of deuterons (# D) and ii) percentage deuteration compared to a fully-deuterated sample (% D). Standard-deviation for each of the triplicate measurements are shown.**

**Supplementary Table S3. Differences in HDX levels for all analysed peptides between PvRBP2b alone and in complex with PvRBP2b human mAbs. Values are expressed as number of deuterons incorporated.**

**Supplementary Table S4. Differences in HDX levels for all analysed peptides between PvRBP2b alone and in complex with PvRBP2b human mAbs. Values are expressed as percentage peptide deuteration.**

**Supplementary Table S5. Data collection and refinement statistics for PvRBP2b complexes with PvRBP2b human Fab fragments.**

**Supplementary Table S6. Summary of interactions between PvRBP2b and PvRBP2b human Fab fragments.**
