## Supplementary Table 5 for "Naturally acquired blocking human monoclonal antibodies to *Plasmodium vivax* reticulocyte binding protein 2b"

**Supplementary Table S5 | Data collection and refinement statistics for PvRBP2b complexes with PvRBP2b human Fab fragments.**

| | PvRBP2b-237235 (PDB 6WM9) | PvRBP2b-241242 (PDB 6WN1) | PvRBP2b-243244 (PDB 6WNO) | PvRBP2b-251249 (PDB 6WOZ) | PvRBP2b-253245 (PDB 6WTY) |
|---|---|---|---|---|---|
| **Data collection*** | | | | | |
| Space group | P 1 | C 1 2 1 | P 1 | P 1 $2_1$ 1 | P 1 $2_1$ 1 |
| Cell dimensions | | | | | |
| *a*, *b*, *c* (Å) | 61.90, 86.78, 90.77 | 360.70, 43.70, 115.33 | 61.01, 84.44, 90.91 | 99.57, 163.55, 121.76 | 70.56, 78.19, 312.08 |
| $\alpha, \beta, \gamma$ (°) | 91.62, 109.98, 99.88 | 90.00, 101.69, 90.00 | 90.73, 110.12, 103.10 | 90.00, 99.19, 90.00 | 90.00, 94.16, 90.00 |
| Resolution (Å) | 43.67-2.45 (2.51-2.45) | 43.72-3.15 (3.23-3.15) | 43.00-3.35 (3.44-3.35) | 49.14-2.90 (3.07-2.90) | 48.70-3.48 (3.69-3.48) |
| $R_{meas}$ | 17.9 (73.6) | 17.0 (121.5) | 21.6 (98.8) | 19.5 (148.4) | 85.7 (317.9) |
| $I/\sigma(I)$ | 7.4 (1.8) | 8.87 (1.32) | 6.9 (1.6) | 9.8 (1.3) | 2.54 (0.53) |
| $CC_{1/2}$ (%) | 98.8 (72.4) | 99.4 (57.2) | 98.9 (68.5) | 99.7 (65.3) | 85.0 (13.1) |
| Completeness (%) | 97.4 (96.6) | 99.6 (99.9) | 97.7 (97.4) | 99.4 (96.6) | 98.7 (93.1) |
| Redundancy | 3.3 (3.4) | 3.79 (3.76) | 3.5 (3.5) | 7.08 (6.72) | 5.4 (5.1) |
| Wilson *B* (Å$^2$) | 55.2 | 64.5 | 68.2 | 61.0 | 60.6 |
| **Refinement** | | | | | |
| No. reflections | 62,347 | 31,277 | 23,187 | 84,919 | 43,312 |
| $R_{work}$ / $R_{free}$ (%) | 22.32 / 27.88 | 25.05 / 28.32 | 24.46 / 28.35 | 21.11 / 25.77 | 28.06/32.40 |
| No. atoms | | | | | |
| Protein | 11,251 | 8,910 | 10,596 | 22,580 | 19,928 |
| Water | 559 | - | - | - | - |
| *B* factors | | | | | |
| Protein | 33.6 | 76.5 | 75.0 | 61.8 | 69.3 |
| Water | 32.0 | - | - | - | - |
| R.m.s. deviations | | | | | |
| Bond lengths (Å) | 0.002 | 0.002 | 0.002 | 0.002 | 0.002 |
| Bond angles (°) | 0.489 | 0.441 | 0.456 | 0.482 | 0.446 |
| Validation | | | | | |
| MolProbity score | 1.36 | 1.58 | 1.65 | 1.45 | 1.49 |
| Clashscore | 4.88 | 5.84 | 7.40 | 4.19 | 4.20 |
| Poor rotamers (%) | 0 | 0 | 0 | 0 | 0 |
| Ramachandran plot | | | | | |
| Favored (%) | 97.45 | 96.16 | 96.35 | 96.27 | 95.89 |
| Allowed (%) | 2.55 | 3.84 | 3.65 | 3.73 | 4.11 |
| Disallowed (%) | 0 | 0 | 0 | 0 | 0 |

**Table S2 | continued**

| | PvRBP2b-258259 (PDB 6WTV) | PvRBP2b-273264 (PDB 6WTU) | PvRBP2b-283284 (PDB 6WQO) |
|---|---|---|---|
| **Data collection*** | | | |
| Space group | P 1 | P 1 | P $2_1$ $2_1$ $2_1$ |
| Cell dimensions | | | |
| *a*, *b*, *c* (Å) | 93.80, 98.65, 103.09 | 93.99, 99.51, 103.49 | 88.25, 126.18, 149.63 |
| *α*, *β*, *γ* (°) | 114.15, 105.00, 89.69 | 114.72, 104.98, 89.81 | 90.00, 90.00, 90.00 |
| Resolution (Å) | 45.01-3.05 (3.23-3.05) | 49.03-2.55 (2.62-2.55) | 42.32-3.15 (3.23-3.15) |
| $R_{meas}$ | 22.6 (65.5) | 10.7 (94.3) | 27.8(129.7) |
| $I/\sigma$ ($I$) | 5.16 (1.97) | 10.01 (1.53) | 6.25 (1.43) |
| $CC_{1/2}$ (%) | 97.1 (65.5) | 99.7 (64.5) | 98.2 (47.4) |
| Completeness (%) | 97.4 (94.9) | 98.2 (97.9) | 99.3 (97.7) |
| Redundancy | 2.5 (2.5) | 3.6 (3.6) | 5.5 (5.7) |
| Wilson *B* ($Å^2$) | 37.8 | 54.6 | 52.2 |
| **Refinement** | | | |
| No. reflections | 60,481 | 104,773 | 29,354 |
| $R_{work}$ / $R_{free}$ (%) | 23.52 / 27.67 | 20.82 / 25.47 | 25.42 / 30.48 |
| No. atoms | | | |
| Protein | 20,978 | 22,128 | 10,482 |
| Water | - | 392 | - |
| *B* factors | | | |
| Protein | 40.5 | 55.7 | 62.3 |
| Water | - | 46.7 | - |
| R.m.s. deviations | | | |
| Bond lengths (Å) | 0.002 | 0.002 | 0.002 |
| Bond angles (°) | 0.436 | 0.497 | 0.467 |
| Validation | | | |
| MolProbity score | 1.65 | 1.44 | 1.68 |
| Clashscore | 6.34 | 4.33 | 6.62 |
| Poor rotamers (%) | 0 | 0 | 0 |
| Ramachandran plot | | | |
| Favored (%) | 95.71 | 96.51 | 95.54 |
| Allowed (%) | 4.29 | 3.49 | 4.46 |
| Disallowed (%) | 0 | 0 | 0 |

X-ray diffraction data were collected on single crystals.
* Values in parentheses are for highest-resolution shell.
