## Supplementary Table 6 for "Naturally acquired blocking human monoclonal antibodies to *Plasmodium vivax* reticulocyte binding protein 2b"

**Supplementary Table S6 | Summary of interactions between PvRBP2b and PvRBP2b human Fab fragments.**

PvRBP2b and 237235 Fab fragment based on the crystal structure PDB 6WM9

| PvRBP2b | Group | Location | 237235 $V_H$ | Group | Distance (Å) |
|---|---|---|---|---|---|
| Hydrogen bonds | | | | | |
| Asn 174 | ND2 | N-ter | Asn 106 | O | 3.0 |
| Arg 193 | NH2 | β2–α1 | Glu 109 | O | 3.5 |
| Arg 193 | N | β2–α1 | Glu 109 | OE1 | 3.1 |
| Arg 193 | NH2 | β2–α1 | Asn 110 | OD1 | 3.0 |
| Arg 193 | NH1 | β2–α1 | Asn 110 | OD1 | 3.0 |
| Lys 437 | NZ | α7 | Tyr 58 | OH | 3.1 |
| Lys 437 | NZ | α7 | Ser 61 | OG | 3.8 |
| Asp 438 | OD1 | α7 | Tyr 58 | OH | 2.6 |
| Asn 441 | ND2 | α7 | Ser 36 | O | 3.1 |
| Lys 445 | NZ | α7 | Ser 36 | O | 2.8 |
| Salt bridges | | | | | |
| Arg 193 | NE | β2–α1 | Glu 109 | OE1 | 3.8 |
| Arg 193 | NE | β2–α1 | Glu 109 | OE2 | 4.0 |
| Lys 434 | NZ | α7 | Asp 62 | OD2 | 2.8 |
| Lys 437 | NZ | α7 | Asp 59 | OD1 | 3.7 |
| Lys 437 | NZ | α7 | Asp 59 | OD2 | 3.1 |

| PvRBP2b | Group | Location | 237235 $V_L$ | Group | Distance (Å) |
|---|---|---|---|---|---|
| Hydrogen bonds | | | | | |
| Asn 231 | OD1 | α2 | Thr 101 | N | 2.9 |
| Asn 231 | OD1 | α2 | Thr 101 | OG1 | 3.8 |
| Asn 231 | ND2 | α2 | Thr 101 | OG1 | 3.5 |
| Arg 235 | NH1 | α2 | Phe 36 | O | 3.2 |
| Gly 238 | O | α2 | Asn 38 | ND2 | 2.9 |
| Asn 241 | ND2 | α2 | Asn 38 | O | 3.2 |

| Other PvRBP2b interfacing residues (237235 $V_H$) | | | | | |
|---|---|---|---|---|---|
| Asp 173 | Asp 176 | Gln 191 | Leu 192 | His 195 | Tyr 200 |
| Asp 203 | Arg 207 | Lys 244 | Lys 248 | Lys 334 | Asn 444 |
| Ile 448 | His 449 | Lys 452 | | | |

| Other PvRBP2b interfacing residues (237235 $V_L$) | | | | | |
|---|---|---|---|---|---|
| Thr 171 | Asp 173 | Arg 193 | Pro 194 | Ile 199 | Tyr 200 |
| Asp 203 | Asn 227 | Leu 230 | Glu 232 | Ser 234 | Ile 237 |
| Ile 239 | Lys 244 | Asn 245 | Lys 248 | | |

PvRBP2b and 241242 Fab fragment based on the crystal structure PDB 6WN1

| PvRBP2b | Group | Location | 241242 $V_H$ | Group | Distance (Å) |
|---|---|---|---|---|---|
| Hydrogen bonds | | | | | |
| Ser 181 | O | β1 | Arg 35 | NH1 | 3.1 |
| Asp 182 | O | β1 | Arg 35 | NH2 | 2.3 |
| Lys 278 | NZ | α3 | Tyr 108 | OH | 3.1 |
| Glu 279 | OE2 | α3 | Asn 36 | ND2 | 3.8 |
| Tyr 351 | OH | α5 | Tyr 111 | OH | 3.0 |
| Lys 379 | O | α6 | Ser 107 | OG | 3.6 |
| Gly 382 | O | α6 | Thr 109 | N | 3.6 |
| Gly 382 | O | α6 | Thr 109 | OG1 | 3.4 |
| Asp 386 | OD1 | α6 | Ala 58 | N | 3.7 |
| Asp 386 | OD2 | α6 | Ser 59 | N | 3.4 |
| Asp 386 | OD1 | α6 | Ser 59 | N | 3.4 |
| Asp 386 | O | α6 | Ser 59 | OG | 3.3 |
| Asp 386 | OD1 | α6 | Ser 59 | OG | 2.8 |
| Glu 389 | OE1 | α6 | Ser 57 | OG | 3.3 |
| Glu 389 | OE1 | α6 | Ala 62 | N | 3.8 |

| PvRBP2b | Group | Location | 241242 $V_L$ | Group | Distance (Å) |
|---|---|---|---|---|---|
| Hydrogen bonds | | | | | |
| Lys 360 | NZ | α5 | Lys 35 | O | 2.5 |
| Lys 363 | NZ | α5 | Tyr 54 | O | 2.5 |
| Salt bridges | | | | | |
| Asp 356 | OD2 | α5 | Arg 95 | NE | 3 |
| Lys 360 | NZ | α5 | Asp 55 | OD1 | 3.5 |
| Lys 360 | NZ | α5 | Asp 55 | OD2 | 2.6 |

| Other PvRBP2b interfacing residues (241242 $V_H$) | | | | | |
|---|---|---|---|---|---|
| Ile 180 | Glu 183 | Ser 184 | Asn 185 | Tyr 186 | Glu 275 |
| Arg 282 | Lys 348 | Leu 352 | Val 355 | Asp 356 | Arg 359 |
| Lys 375 | Gln 378 | Tyr 383 | Ile 385 | Arg 387 | Tyr 390 |
| Gln 393 | | | | | |

| Other PvRBP2b interfacing residues (241242 $V_L$) | | | | | |
|---|---|---|---|---|---|
| Lys 348 | Asn 349 | Leu 352 | Ser 353 | Val 355 | Arg 359 |
| Glu 362 | | | | | |

PvRBP2b and 243244 Fab fragment based on the crystal structure PDB 6WNO

| PvRBP2b | Group | Location | 243244 $V_H$ | Group | Distance (Å) |
|---|---|---|---|---|---|
| Hydrogen bonds | | | | | |
| Arg 193 | N | β2–α1 | Glu 109 | OE2 | 2.8 |
| Arg 193 | NH2 | β2–α1 | Asn 110 | OD1 | 2.7 |
| Tyr 200 | OH | α1 | Asn 62 | ND2 | 3.8 |
| Lys 437 | O | α7 | Tyr 58 | OH | 3.2 |
| Asp 438 | OD1 | α7 | Tyr 58 | OH | 2.4 |
| Asn 441 | ND2 | α7 | Ser 36 | O | 3.1 |
| Lys 445 | NZ | α7 | Ser 36 | O | 3.5 |
| Salt bridges | | | | | |
| Arg 193 | NH2 | β2–α1 | Glu 109 | OE2 | 3.9 |
| Lys 437 | NZ | α7 | Asp 59 | OD2 | 3.7 |
| Lys 437 | NZ | α7 | Asp 61 | OD2 | 3.5 |

| PvRBP2b | Group | Location | 243244 $V_L$ | Group | Distance (Å) |
|---|---|---|---|---|---|
| Hydrogen bonds | | | | | |
| Asn 231 | OD1 | α2 | Thr 101 | N | 3.1 |
| Arg 235 | NH1 | α2 | Phe 36 | O | 3.5 |
| Asn 241 | ND2 | α2 | Asn 38 | O | 3.1 |

| Other PvRBP2b interfacing residues (243244 $V_H$) | | | | | |
|---|---|---|---|---|---|
| Asp 173 | Asn 174 | Asp 176 | Gln 191 | Leu 192 | HIS 195 |
| Asp 203 | Lys 334 | Tyr 430 | | | |

| Other PvRBP2b interfacing residues (243244 $V_L$) | | | | | |
|---|---|---|---|---|---|
| Thr 171 | Asp 173 | Arg 193 | Pro 194 | HIS 195 | Ile 199 |
| Tyr 200 | Asp 203 | Asn 227 | Leu 230 | Glu 232 | Ser 234 |
| Ile 237 | Gly 238 | Ile 239 | Lys 244 | Asn 245 | |

### PvRBP2b and 251249 Fab fragment based on the crystal structure PDB 6WOZ

| PvRBP2b | Group | Location | 251249 $V_H$ | Group | Distance (Å) |
|---|---|---|---|---|---|
| Hydrogen bonds | | | | | |
| Tyr 186 | OH | β1–β2 | Gly 109 | N | 3.6 |
| Tyr 186 | OH | β1–β2 | Gly 109 | O | 2.8 |
| Arg 290 | NH1 | α3 | Tyr 59 | OH | 3.5 |
| Lys 291 | NZ | α3 | Tyr 59 | O | 3.3 |
| Lys 291 | NZ | α3 | Ser 60 | O | 3.7 |
| Asp 294 | OD2 | α3 | Thr 57 | OG1 | 2.7 |
| Asp 294 | OD2 | α3 | Gly 58 | N | 3.4 |
| Asp 294 | OD2 | α3 | Tyr 59 | N | 2.7 |
| Asp 294 | OD2 | α3 | Ser 60 | N | 3.7 |
| Asp 294 | OD2 | α3 | Ser 61 | N | 3.4 |
| Gln 295 | NE2 | α3 | Ser 61 | OG | 2.8 |
| Gln 295 | NE2 | α3 | Glu 62 | OE2 | 3.7 |
| Lys 297 | NZ | α3 | Ser 36 | O | 3.3 |
| Lys 297 | NZ | α3 | Ser 38 | OG | 3.8 |
| Asn 300 | ND2 | α3 | Ile 112 | O | 3.2 |
| Asn 301 | OD1 | α3 | Asp 114 | N | 2.6 |
| Arg 304 | NH1 | α3 | Asp 114 | O | 2.8 |
| Asn 428 | ND2 | α7 | Val 110 | O | 3.7 |
| Salt bridges | | | | | |
| Lys 298 | NZ | α3 | Glu 62 | OE1 | 3.7 |
| Lys 298 | NZ | α3 | Glu 62 | OE2 | 2.6 |

| PvRBP2b | Group | Location | 251249 $V_L$ | Group | Distance (Å) |
|---|---|---|---|---|---|
| Hydrogen bonds | | | | | |
| Glu 308 | OE2 | α3 | Thr 36 | OG1 | 3.1 |
| Salt bridges | | | | | |
| Asp 305 | OD1 | α3 | Arg 96 | NH1 | 3.6 |
| Asp 305 | OD1 | α3 | Arg 96 | NH2 | 2.9 |
| Asp 305 | OD2 | α3 | Arg 96 | NH1 | 3.3 |
| Other PvRBP2b interfacing residues (251249 $V_H$) | | | | | |
| Asp 182 | Glu 183 | Ser 184 | Tyr 187 | Asn 198 | PHE 201 |
| Leu 293 | Glu 421 | PHE 424 | Asn 425 | Tyr 429 | |
| Other PvRBP2b interfacing residues (251249 $V_L$) | | | | | |
| Arg 304 | Asn 417 | | | | |

### PvRBP2b and 253245 Fab fragment based on the crystal structure PDB 6WTY*

| PvRBP2b | Group | Location | 253245 $V_H$ | Group | Distance (Å) |
|---|---|---|---|---|---|
| Hydrogen bonds | | | | | |
| Gln 393 | NE2 | | Ser 112 | O | 3.0 |
| Salt Bridges | | | | | |
| Asp 386 | OD1 | α6 | Arg 114 | NH2 | 3.6 |
| Glu 389 | OE1 | α6 | Arg 114 | NH2 | 2.5 |
| Glu 389 | OE2 | α6 | Arg 114 | NH2 | 3.4 |

| PvRBP2b | Group | Location | 253245 $V_L$ | Group | Distance (Å) |
|---|---|---|---|---|---|
| Hydrogen bonds | | | | | |
| Tyr 429 | OH | α7 | Gly 35 | O | 3.8 |
| Other PvRBP2b interfacing residues (253245 $V_H$) | | | | | |
| Asp 341 | Ile 344 | Lys 345 | Lys 348 | Asn 349 | Tyr 351 |
| Leu 352 | Val 355 | Asp 356 | Lys 379 | Gly 382 | Ile 385 |
| Leu 392 | | | | | |
| Other PvRBP2b interfacing residues (253245 $V_L$) | | | | | |
| Ser 184 | Asn 185 | Tyr 186 | Asp 386 | Arg 387 | Tyr 390 |
| Gln 393 | Lys 394 | Lys 396 | His 397 | Asp 400 | Gln 401 |
| Ala 404 | Tyr 422 | Asn 425 | Asn 428 | | |

### PvRBP2b and 258259 Fab fragment based on the crystal structure PDB 6WTV

| PvRBP2b | Group | Location | 258259 $V_H$ | Group | Distance (Å) |
|---|---|---|---|---|---|
| Hydrogen bonds | | | | | |
| Glu 183 | O | β1–β2 | Ser 80 | OG | 3.1 |
| Asn 185 | N | β1–β2 | Ser 80 | OG | 3.8 |
| Lys 348 | NZ | α5 | Ser 62 | OG | 3.2 |
| Lys 348 | NZ | α5 | Tyr 64 | OH | 2.3 |
| Tyr 351 | OH | α5 | Tyr 108 | O | 2.7 |
| Arg 359 | NH1 | α5 | HIS 112 | O | 2.5 |
| Asp 386 | OD1 | α6 | Gly 58 | N | 3.6 |
| Asp 386 | OD1 | α6 | Ser 59 | N | 3.1 |
| Asp 386 | OD2 | α6 | Ser 59 | N | 3.2 |
| Asp 386 | O | α6 | Ser 59 | OG | 2.5 |
| Asp 386 | OD1 | α6 | Ser 59 | OG | 2.6 |
| Salt bridges | | | | | |
| Asp 356 | OD1 | α5 | HIS 112 | NE2 | 3.5 |
| Asp 356 | OD2 | α5 | HIS 112 | NE2 | 3.6 |
| Lys 379 | NZ | α6 | Glu 107 | OE2 | 3.9 |

| PvRBP2b | Group | Location | 258259 $V_L$ | Group | Distance (Å) |
|---|---|---|---|---|---|
| Hydrogen bonds | | | | | |
| Asp 356 | OD1 | α5 | Thr 36 | OG1 | 3.1 |
| Asp 356 | OD2 | α5 | Thr 36 | N | 3.7 |
| Other PvRBP2b interfacing residues (258259 $V_H$) | | | | | |
| Ser 184 | Lys 278 | Leu 352 | Ser 353 | Val 355 | Gly 382 |
| Tyr 383 | Ile 385 | Arg 387 | Glu 389 | Tyr 390 | Gln 393 |
| Other PvRBP2b interfacing residues (258259 $V_L$) | | | | | |
| Lys 348 | Asn 349 | Leu 352 | Arg 359 | Lys 360 | |

PvRBP2b and 273264 Fab fragment based on the crystal structure PDB 6WTU

| PvRBP2b | Group | Location | 273264 $V_H$ | Group | Distance (Å) |
|---|---|---|---|---|---|
| Hydrogen bonds | | | | | |
| Tyr 351 | OH | α5 | Tyr 108 | O | 2.6 |
| Arg 359 | NH2 | α5 | HIS 112 | O | 2.5 |
| Asp 386 | OD1 | α6 | Gly 58 | N | 3.8 |
| Asp 386 | OD1 | α6 | Ser 59 | N | 3.3 |
| Asp 386 | OD2 | α6 | Ser 59 | N | 3.5 |
| Asp 386 | OD1 | α6 | Ser 59 | OG | 3.1 |
| Asp 386 | O | α6 | Ser 59 | OG | 3.0 |
| Glu 389 | OE1 | α6 | Ser 62 | OG | 2.7 |
| Gln 393 | NE2 | α6 | Gly 60 | O | 3.4 |
| Salt bridges | | | | | |
| Asp 356 | OD2 | α5 | HIS 112 | NE2 | 3.8 |

| PvRBP2b | Group | Location | 273264 $V_L$ | Group | Distance (Å) |
|---|---|---|---|---|---|
| Hydrogen bonds | | | | | |
| Asp 356 | OD2 | α5 | Thr 35 | OG1 | 3.0 |
| Asp 356 | OD1 | α5 | Ser 36 | N | 3.6 |
| Asp 356 | OD1 | α5 | Ser 36 | OG | 2.4 |
| Asp 356 | OD2 | α5 | Ser 36 | N | 3.4 |
| Lys 360 | NZ | α5 | Thr 35 | O | 2.6 |
| Salt bridges | | | | | |
| Lys 360 | NZ | α5 | Asp 55 | OD1 | 3.5 |
| Lys 360 | NZ | α5 | Asp 55 | OD2 | 2.5 |
| Other PvRBP2b interfacing residues (273264 $V_H$) | | | | | |
| Glu 183 | Ser 184 | Asn 185 | Lys 348 | Leu 352 | Ser 353 |
| Val 355 | Lys 379 | Gly 382 | Tyr 383 | Ile 385 | Arg 387 |
| Tyr 390 | | | | | |
| Other PvRBP2b interfacing residues (273264 $V_L$) | | | | | |
| Asn 349 | Leu 352 | Arg 359 | | | |

PvRBP2b and 283284 Fab fragment based on the crystal structure PDB 6WQO

| PvRBP2b | Group | Location | 283284 $V_H$ | Group | Distance (Å) |
|---|---|---|---|---|---|
| Hydrogen bonds | | | | | |
| Tyr 211 | O | α1 | Ser 35 | OG | 2.3 |
| Tyr 211 | OH | α1 | Ile 104 | O | 3.2 |
| Asn 319 | ND2 | α4 | Tyr 37 | OH | 3.6 |
| Asp 323 | OD2 | α4 | Tyr 37 | OH | 2.8 |
| Asp 331 | OD1 | α4 | Tyr 108 | OH | 2.5 |
| Salt bridges | | | | | |
| Lys 216 | NZ | α2 | Asp 58 | OD2 | 2.4 |
| Asp 323 | OD2 | α4 | Lys 102 | NZ | 2.8 |
| Asp 323 | OD1 | α4 | Lys 102 | NZ | 4.0 |

| PvRBP2b | Group | Location | 283284 $V_L$ | Group | Distance (Å) |
|---|---|---|---|---|---|
| Hydrogen bonds | | | | | |
| Lys 326 | NZ | α4 | Ser 61 | OG | 3.8 |
| Lys 333 | NZ | α4 | Asp 55 | O | 3.1 |
| Salt bridges | | | | | |
| Lys 326 | NZ | α4 | Glu 60 | OE1 | 3.7 |
| Lys 326 | NZ | α4 | Glu 60 | OE2 | 3.5 |
| Lys 333 | NZ | α4 | Asp 55 | OD2 | 2.5 |
| Other PvRBP2b interfacing residues (283284 $V_H$) | | | | | |
| Arg 207 | Ser 210 | HIS 212 | Thr 213 | Gln 317 | Val 320 |
| Met 324 | Lys 326 | Ile 327 | Val 330 | Lys 412 | |
| Other PvRBP2b interfacing residues (283284 $V_L$) | | | | | |
| Val 330 | Lys 334 | Lys 410 | | | |

The distance measurements are based on molecules A, B and C.
Interacting and interfacing residues between PvRBP2b and antibody Fabs was determined using PISA[31].
*Some residue side chains are unresolved and the interactions listed here may not be complete.
